# *prdm1a* drives a fate switch between hair cells of different mechanosensory organs

**DOI:** 10.1101/2025.06.01.657251

**Authors:** Jeremy E Sandler, Ya-Yin Tsai, Shiyuan Chen, Logan Sabin, Mark E. Lush, Abhinav Sur, Elizabeth Ellis, Nhung TT Tran, Malcolm Cook, Allison R Scott, Jonathan S. Kniss, Jeffrey A. Farrell, Tatjana Piotrowski

## Abstract

Vertebrate mechanosensory hair cells (HCs) in the ear detect sound and gravitational forces. Additionally, fish have homologous lateral line HCs in the skin that detect water vibrations for orientation and predator avoidance. HCs in fish and other non-mammalian vertebrates regenerate to restore function after damage, but mammalian HCs lack this ability, causing deafness and vestibular defects. Experimental attempts at regeneration in mice result in incomplete differentiation of immature HCs. Despite differences in regeneration, the gene regulatory networks (GRNs) driving HC maturation are highly similar across vertebrates. Here, we show that the transcription factor *prdm1a* plays a key role in the HC fate GRN in the zebrafish lateral line. Mutating *prdm1a* respecifies lateral line HCs into ear HCs, altering morphology and transcriptome. Understanding how transcription factors control diverse HC fates in zebrafish is crucial for understanding the yet unsolved regeneration of diverse HCs in mammalian ears to restore hearing and balance.

## Introduction

Vertebrates possess mechanosensory hair cells in the ear that transduce vibrations into nerve impulses interpreted as sound.^1, 2^ Fishes have two separate sensory organ systems with functionally and genetically similar mechanosensory cells; the ear responsible for hearing and orientation, and the lateral line responsible for sensing water vibrations and detecting external physical stimuli (Figs. 1a-1c).^3^ Within the ear, there are several sensory epithelia containing diverse hair cells with specialized functions.^4, 5, 6^ In the mammalian cochlea, inner hair cells (IHCs) and outer hair cells (OHCs) are responsible for hearing,^1^ while Type I and Type II hair cells in the saccule and utricle detect linear acceleration, and the cristae of the semicircular canals detect rotational acceleration. In the saccule and utricle the two types of hair cells are divided into striolar and extra-striolar regions based on polarity and location within the organs.^7^ Early developmental specification of these different hair cell types is controlled by common gene regulatory networks (GRNs),^8, 9, 10^ but control of terminal differentiation is exerted by distinct GRNs that result in different functions and physical characteristics.^11, 12, 13^ The precise specification of each hair cell type in the ear is essential for hearing and vestibular function, and incorrect developmental fate leads to profound hearing loss.^1, 14, 15, 16^

**Fig. 1.**
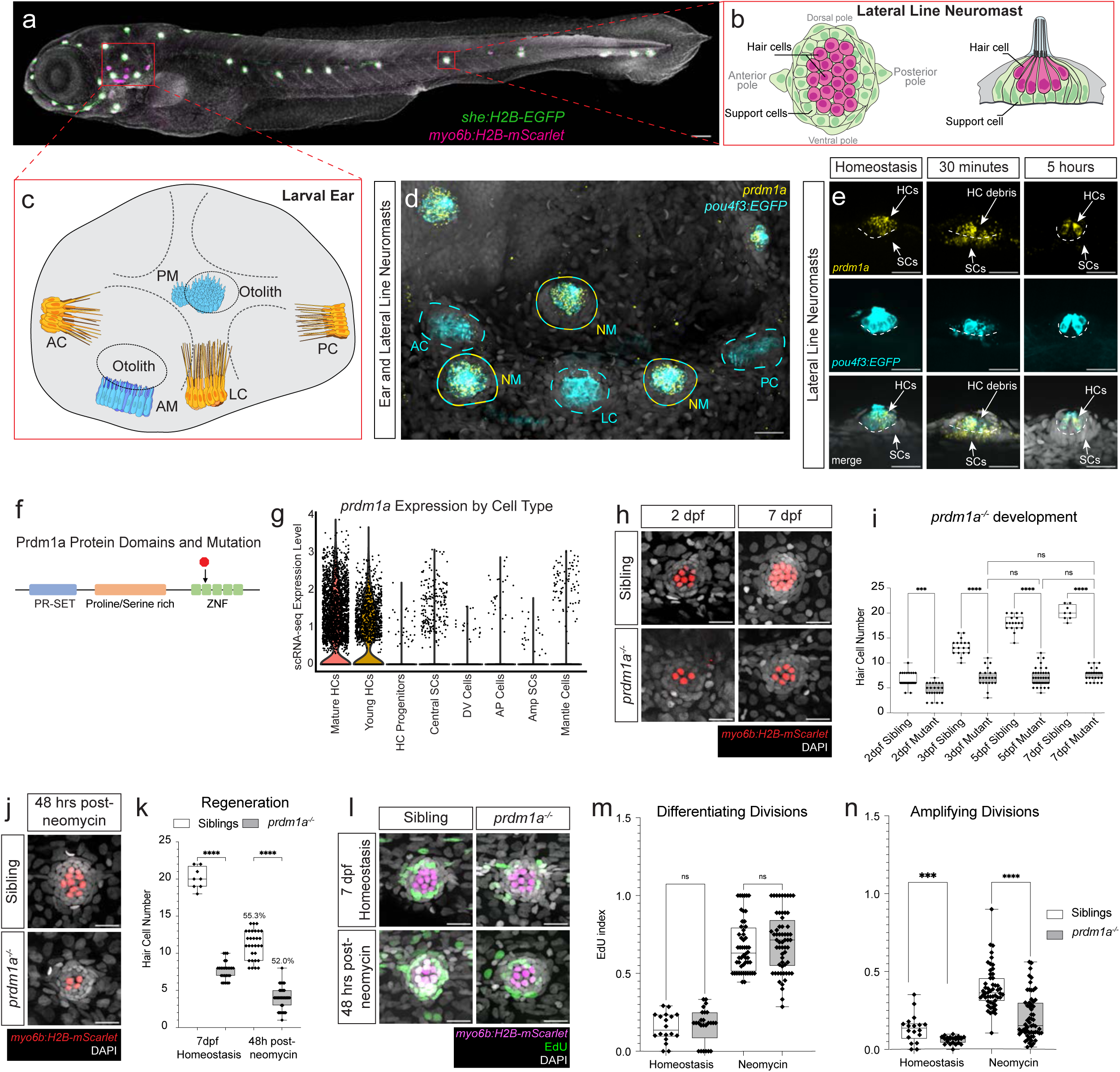
*prdm1a* expression is integral to hair cells with mutant phenotypes during development and regeneration. (a) A 5 dpf zebrafish embryo with hair cells expressing *Tg(myo6b:H2B-mScarlet)* in magenta and support cells expressing *Tg(she:H2B-EGFP)* in green, respectively. Scale bar= 100 µm. (b) Schematic of the lateral line with hair cells and support cells in magenta and green, respectively. (c) Schematic of the zebrafish ear. AC: anterior crista, LC: lateral crista, PC: posterior crista, AM: anterior macula, PM: posterior macula. (d) HCR of lateral line neuromasts and cristae of the zebrafish ear with *prdm1a* probe and *Tg(pou4f3:GAP-EGFP)* in hair cells. (e) *prdm1a* expression in the lateral line during homeostasis, and during regeneration at 30 minutes and 5 hours post neomycin, with *Tg(pou4f3:GAP-EGFP)* in hair cells. HC: Hair Cells, SC: Support Cells. (f) Schematic of Prdm1a protein with major domains and stop codon in *prdm1a* mutant (black arrow). (g) Violin plot of *prdm1a* expression from sibling scRNA-seq, divided by cell type. (h) Hair cells labeled with *Tg(myo6b:H2B-mScarlet)* show developmental defects in *prdm1a* mutant neuromasts at 2 and 7 dpf compared to siblings. (i) Quantification of hair cells during development in sibling and *prdm1a^-/-^* neuromasts. n=6 fish per timepoint and genotype, 3-4 neuromasts per fish. Two-way Anova with Tukey’s multiple comparisons test, **** p<0.0001. (j) Hair cells labeled with *Tg(myo6b:H2B-mScarlet)* show regeneration defects in *prdm1a^-/-^* neuromasts 48 hours after neomycin treatment; 7 dpf. (k) Quantification of hair cells during regeneration and homeostasis at the same developmental timepoint, in sibling and *prdm1a^-/-^* neuromasts. The percentage of regenerated hair cells compared to homeostasis for each condition is shown above regeneration counts. n= 6 fish per timepoint and genotype, 3-4 neuromasts per fish. Two-way Anova with Tukey’s multiple comparisons test, **** p<0.0001. (l) EdU (green) incorporation and hair cells expressing *Tg(myo6b:H2B-mScarlet)* (magenta) during development and regeneration in siblings and mutants. (m and n) Quantification of EdU incorporation in differentiating divisions (m) and amplifying divisions (n) during regeneration in sibling and *prdm1a^-/-^*neuromasts. n= 6-20 fish per treatment and genotype, 3-4 neuromasts per fish. Student’s t-test, **** p<0.0001. Scale bars = 20 µm.

The ears of aquatic vertebrates also have a similar diversity of hair cell types and functions (Figs 1a and 1c).^17^ Like mammals, adult fishes possess vestibular hair cells in cristae, utricles, saccules and the later-developing lagena. Fishes do not possess a cochlea, but the utricle, saccule and lagena also serve an auditory function.^17^ Fish lateral line hair cells share genetic, functional, and structural similarity with ear hair cells of many species,^6, 18^ including core gene networks that regulate their early specification.^8, 19, 20^ Likewise, genes linked to deafness in mammals cause defects in hair cell function when mutated in zebrafish.^21, 22, 23^

Despite these similarities, there is a striking disparity of hair cell regeneration between fish and mammals. While hair cells in non-mammalian vertebrates regenerate following death,^6, 24, 25^ mammalian hair cells lack this ability,^9, 26, 27^ causing hearing loss and vestibular dysfunction in humans.^28, 29^ During development and regeneration of the zebrafish lateral line, hair cells arise from central support cells that turn into hair cell progenitors, then divide and form young hair cells before differentiating into mature hair cells (Fig. 1b).^19, 30^ Attempts at stimulating hair cell regeneration in mammalian cochleae through manipulating signaling pathways or gene therapy result in incomplete differentiation or hair cells of only one type.^31, 32, 33, 34, 35, 36, 37, 38, 39, 40^ Therefore, understanding the unique developmental requirements for each hair cell type is prerequisite to any attempt to restore proper function. As mammals lack the molecular and genetic programs to regenerate hair cells, studying the process in a robustly regenerating model provides a powerful tool to identify missing components and gain insights for broader applications.

We recently performed a single-cell RNA sequencing (scRNA-seq) time course of lateral line hair cell regeneration, and identified genes and pathways necessary for proliferation and regeneration.^19^ In addition, we and others have characterized single cell transcriptomic profiles of the zebrafish lateral line and ear during homeostasis and development to provide a comprehensive transcriptional atlas.^5, 20, 41^ We searched these datasets for genes possibly involved in regeneration and identified the transcription factor *prdm1a*, which is expressed both during hair cell differentiation and regeneration in the lateral line. *prdm1a* is not expressed in ear hair cells,^12^ pointing to a unique role in hair cell specification and regeneration in the lateral line. Prdm1a is a direct and indirect transcriptional repressor, but can also activate certain genes.^42, 43, 44^ In mammals, *Prdm1* (*Blimp1*) directs cell fate decisions in a variety of tissues and organs.^45, 46, 47^ In zebrafish, *prdm1a* is a central regulator of neural crest cell proliferation and fate decisions during neurogenesis and specification of slow twitch muscle fibers.^48, 49, 50^ *prdm1a,* therefore, has hallmarks of a gene that controls cell fate, identity, and downstream functional properties across species^51^ and is a prime candidate for investigations into proliferation and hair cell differentiation and regeneration.

We mutated *prdm1a* and observed profound defects in the development and regeneration of lateral line hair cells, and a decrease in support cell proliferation during regeneration. Using scRNA-seq, we characterized transcriptional changes associated with this mutation, and found evidence of a fate switch from lateral line to an ear-like hair cell state, with de-repression of genes normally expressed only in ear hair cells. Morphological characterization of mutant hair cells also indicates a transformation to an ear hair cell like morphology. We show that the ectopically expressed genes in *prdm1a* mutants possess enhancers and promoters that respond to direct transcriptional repression by Prdm1a. Lastly, we present a GRN model to describe upstream and downstream genetic interactions of *prdm1a* in the lateral line that places the gene into the broader context of hair cell specification and regeneration. This GRN shows striking similarities to the core regulatory code shared between lateral line and ear hair cells of fish and mammals. We show that, even though the hair cell GRN is highly similar between the ear and the lateral line, the lack of expression of *prdm1a* in the ear but continued expression of *prdm1a* in the lateral line leads to the differentiation of lateral line hair cells instead of ear hair cells.

## Results

### *prdm1a* expression is integral to the lateral line and hair cells during development and regeneration

*prdm1a* is expressed in central support cells during the early stages of regeneration and subsequently in differentiating hair cells (Fig. 1g).^19^ To analyze hair cell expression in more detail, we performed scRNA-seq on 5 days post fertilization (dpf) homeostatic *prdm1a* mutants (described below) and siblings (Supplementary Fig. 1g). We confirmed that *prdm1a* partially overlaps with the hair cell specifying gene *atoh1a* and is strongly expressed in young and mature hair cells and sometimes in a few support cells (Supplementary Figs. 1h and 1i). *prdm1a* truncated transcript is also detected in mutant cells, even though it is non-functional. We validated the expression results with *in situ* hybridization chain reaction (HCR) in larvae, showing that *prdm1a* is expressed in all hair cells of the lateral line. *prdm1a*, however, is excluded from hair cells in the ear, which still express other hair cell marker genes such as *pou4f3* (Figs. 1d and 1e). To characterize *prdm1a’s* spatiotemporal expression during regeneration, we killed hair cells using neomycin^52^ and performed HCR. We observed a strong upregulation of *prdm1a* expression in central support cells 30 minutes and 1 hour after killing hair cells (Fig. 1e and Supplementary Fig.1a). Some of these central support cells will proliferate and differentiate into regenerated hair cells.^19, 30^ As regeneration progresses, *prdm1a* expression is once again restricted to newly regenerated hair cells 5 hours after hair cell killing, overlapping with *pou4f3*:*EGFP* expression.

We previously identified a mutation in *prdm1a* in an unpublished chemical mutagenesis screen, allowing us to test its function *in vivo*. The mutation causes a premature stop codon in *prdm1a*, truncating the zinc finger DNA binding domain, rendering the protein unable to bind to DNA and regulate target genes (Fig. 1f, black arrow). Consistent with previous findings, we see defects in slow muscle fibers along the myoseptum and in melanocytes.^48, 53, 54^ The defect in myoseptum formation causes the random or stalled migration of the posterior lateral line primordium, leading to irregularly deposited neuromasts along the trunk (Supplementary Figs. 1d and 1e).^55^ The larvae also have small deformed pectoral fins, but look morphologically relatively normal.

We assayed lateral line hair cell development in *prdm1a* mutants, from 2 dpf through neuromast maturation and homeostasis at 7 dpf. We find a failure of *prdm1a* mutant hair cells to increase in number starting at 2 dpf (Figs. 1h and 1i; Supplementary Fig. 1f). The result of this developmental stalling in hair cell formation is a permanent reduction in the number of hair cells compared to sibling embryos. To evaluate the impact of the *prdm1a* mutation on regenerative ability, we killed the hair cells and assayed regeneration in siblings and mutants 48 hours later. In siblings, we observed an average of 11-12 regenerated hair cells, or 55.3% of homeostasis, and 4 hair cells, or 52.0% of homeostasis on average in mutants (Figs. 1j and 1k). Since *prdm1a* mutant neuromasts possess on average 6 or 7 hair cells before neomycin treatment and the regeneration rates are very similar, we conclude that regeneration of hair cells is unaffected beyond the developmental defects we observed.

Hair cell regeneration in the lateral line also crucially depends on proliferation of two different support cell types. Central support cells undergo differentiating divisions and develop into regenerated hair cells, and peripheral support cells undergo amplifying divisions to replenish the central support cell pool.^19, 30^ As a *prdm1a* mutation decreases proliferation in other zebrafish tissues,^50, 56^ we performed EdU incorporation experiments to assay proliferation in siblings and *prdm1a* mutants. Differentiating divisions result in EdU-positive hair cells, while amplifying divisions result in EdU-positive support cells. During both development and regeneration, the ratio of peripheral, amplifying support cells marked by EdU is significantly lower in the *prdm1a* mutant embryos (Figs. 1l and 1n). This indicates a non-cell autonomous role for *prdm1a* in proliferation, as it is not expressed in these amplifying, peripheral support cells. We did not, however, observe a difference in EdU in hair cells themselves (Figs. 1l and 1m), indicating that differentiating divisions are not affected by loss of *prdm1a*, and that *prdm1a* plays a proliferation-independent role in hair cells, consistent with our observations about similar rates of regeneration overall.

Together, these results point to dual functions for *prdm1a* in the lateral line. *prdm1a* plays an indirect non-cell autonomous role in amplifying support cell proliferation, which likely underlies the defect in the development of a normal number of hair cells. While the role of *prdm1a* in proliferation requires further investigation, in this manuscript we explore the previously unknown cell-autonomous role that *prdm1a* plays in postmitotic hair cells.

### scRNA-seq indicates a cell fate switch in *prdm1a* mutant hair cells

The hair cell-specific and regeneration-responsive expression of *prdm1a* in the lateral line led us to conduct a scRNA-seq experiment with *prdm1a* mutants and siblings to determine what role it might be playing in hair cell differentiation. We FAC-sorted 5 dpf zebrafish embryos using a transgenic line driving mCherry in all lateral line cells (*Tg(she:mCherry))*, separately sorting *prdm1a* mutant and sibling embryos, and performed scRNA-seq using the 10X Chromium platform. The resulting datasets were subsequently integrated for downstream analysis. All expected lateral line cell types were present, including sibling and mutant hair cells along our previously established developmental trajectory^19^ (Supplementary Fig.1g, Supplementary Data 1). We subsetted the integrated hair cell lineage from central support cells to mature hair cells, as these are the cell types that express *prdm1a* in the lateral line. Only after subsetting we observed a split in the hair cell developmental trajectory at the young hair cell stage (Fig. 2a, green cells), with hair cells located in two transcriptionally distinct clusters after the split. Strikingly, when we color the cells by genotype, the *prdm1a* mutant hair cells segregate from sibling hair cells (Fig. 2b). The split in hair cell trajectory occurs when *prdm1a* is first expressed (Fig. 2c, black arrow), suggesting that the activity of *prdm1a* is integral to hair cell fate. We performed a differential gene expression analysis between sibling and mutant hair cell clusters to determine the transcriptional changes driving the separate trajectory and clustering of mutant hair cells and identified 1589 upregulated and 344 downregulated genes (Supplemental Data 2). As *prdm1a* acts mostly as a repressor, we focused on the larger set of upregulated genes as likely targets of direct repression, while acknowledging the possibility of indirect interactions as well. These ectopic genes are upregulated primarily in mutant hair cells after they differentiate (Figs. 2d-2h; Supplementary Figs. 2a-2d). The *prdm1a* mutant hair cells still express canonical hair cell markers, such as *myo6b, pou4f3*, *gfi1aa/ab and atoh1a/b* (Fig. 2d) and they also express genes typical for mature hair cells, such as *fgf10b* and the tip link component *pcdh15a* (Supplementary Figs. 2m and 2n), suggesting that they are not delayed in their development. Many of the upregulated, genes cause auditory dysfunction in mice and/or humans when mutated, pointing to their function in ear hair cells (Mutant phenotypes reviewed in Supplemental Table 1A).^14, 57, 58, 59, 60, 61, 62, 63, 64, 65, 66, 67, 68, 69^. We subsetted scRNA-seq data for all other neuromast cell types, but found no additional separate clustering based on *prdm1a* genotype, supporting a hair cell-specific effect of *prdm1a* on gene regulation.

**Fig. 2.**
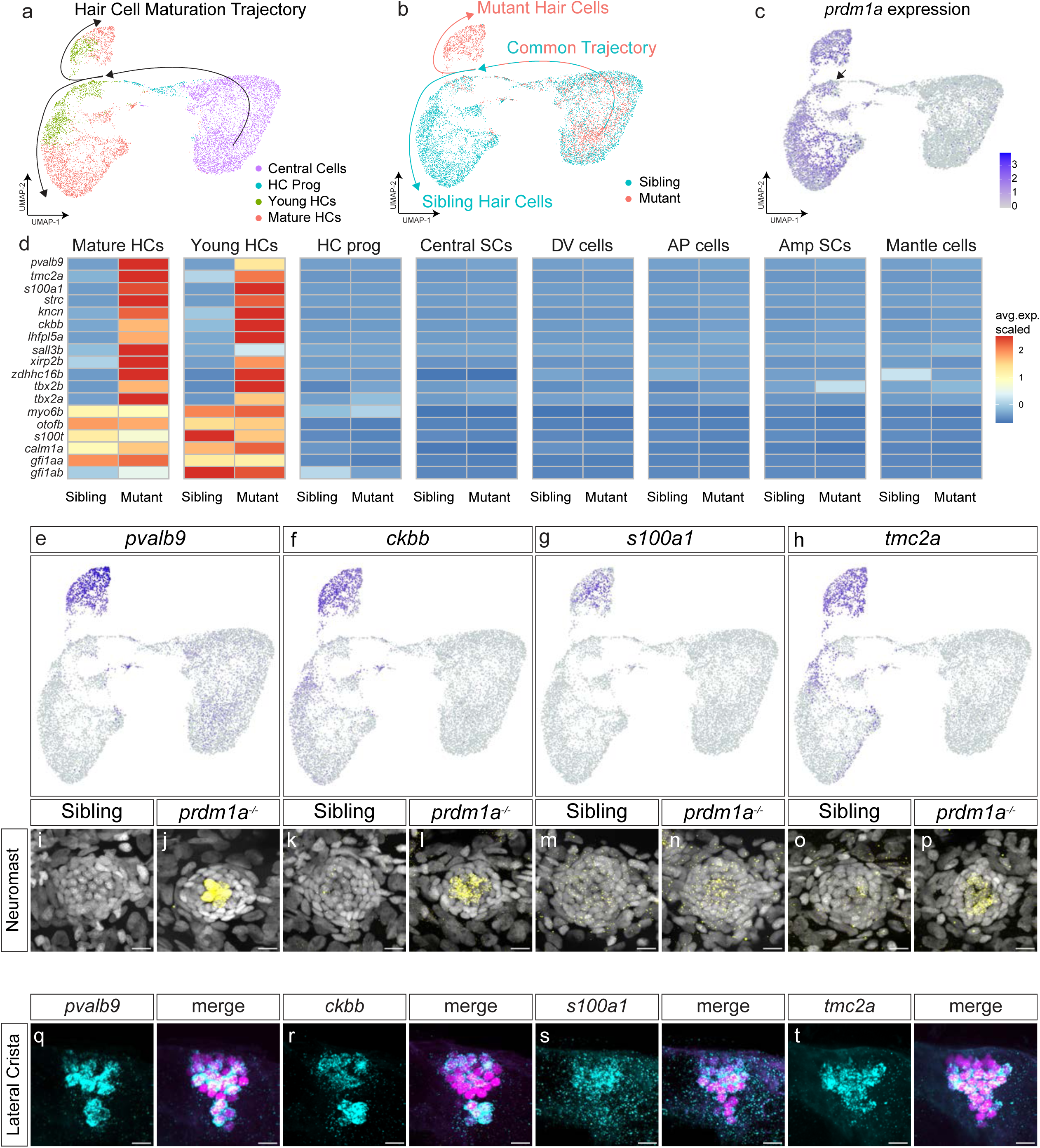
scRNA-seq indicates a hair cell differentiation fate switch in *prdm1a* mutants. (a) UMAP plot of *prdm1a^-/-^* and sibling scRNA-seq, from central support cells to mature hair cells, colored by cell type. Black arrows indicate developmental trajectory. (b) UMAP plot of *prdm1a^-/-^* and sibling scRNA-seq, from central support cells to mature hair cells, colored by genotype. (c) Feature plot of *prdm1a* expression in the scRNA-seq hair cell lineage sub-clustering. (d) Heatmap of selected genes upregulated in *prdm1a^-/-^* hair cells, and non-differentially expressed hair cell marker genes. (e-h) Feature plots of *pvalb9, ckbb, s100a1* and *tmc2a* from the hair cell trajectory sub-clustered scRNA-seq. (i-p) Representative images of HCRs of *pvalb9, ckbb, s100a1* and *tmc2a* in 5 dpf sibling and *prdm1a^-/-^* neuromasts. (q-t) HCRs of *pvalb9, ckbb, s100a1* and *tmc2a* in the lateral crista of the ear in *Tg(myo6b: H2B-mScarlet)* larvae, with hair cells are labeled in magenta. Scale bars= 10 µm.

To validate the expression of genes upregulated in *prdm1a* mutants, we performed HCRs for *pvalb9, ckbb, s100a1, tmc2a*, *strc, kncn, lhfpl5a*, *tbx2a and tbx2b* and found them to all be strongly expressed in the lateral line hair cells of *prdm1a* mutants, but not sibling hair cells (Figs. 2i-2p, Supplementary Figs. 2e-2h). On the other hand, all these genes are strongly expressed in sibling hair cells in the ear (Figs. 2q-2t and Supplementary Figs. 2i-2j). Of particular interest are the family of Tmc genes, which are members of the mechanotransduction complex, as they are differentially expressed in lateral line and ear hair cells^70, 71, 72, 73^. In the sibling lateral line *tmc2a* is only expressed in a few sibling lateral line hair cells but is expressed in all ear (lateral crista) hair cells (Figs. 2h and 2o-2p and 2t).^73^ In *prdm1a* mutants all lateral line hair cells express *tmc2a*, suggesting that mutant lateral line hair cells acquired an ear hair cell fate (Fig. 2p). On the other hand, the paralog *tmc2b* is highly expressed in the sibling and *prdm1a* mutant lateral line hair cells, suggesting that it is not regulated by *prdm1a* (Supplementary Fig. 2l). Also, in the lateral crista in the ear *tmc2b* is only expressed in a few hair cells that do not appear to express *tmc2a* (Supplementary Fig. 2k and Fig. 2t).

The hypothesis that loss of *prdm1a* leads to a fate switch is supported by our analysis of the genes upregulated in *prdm1a* mutant lateral line hair cells in published zebrafish ear and mammalian ear scRNA-seq data.^5, 12, 13, 74, 75^ We used the gEAR data exploration tool^76^ to compile a table of expression of these genes, and show that they are normally expressed in zebrafish and mammalian ear hair cells (Supplementary Table 1). Together, the expression analyses strongly suggest that loss of *prdm1a* causes lateral line hair cells to adopt an ear hair cell fate.

### *prdm1a* regulates the expression of target genes as a transcription factor

*prdm1a* is a strong repressor with a DNA-binding zinc finger domain, and a large number of genes are upregulated specifically in hair cells when it is mutated. Therefore, we asked if Prdm1a regulates these genes by binding to their enhancers and promoters. We performed ATAC-seq on FAC-sorted wildtype lateral line cells during homeostasis. We then performed a motif enrichment analysis specifically on the putative regulatory regions and promoters of genes upregulated in the *prdm1a* mutant hair cells. We find that the Prdm1 binding site is highly enriched in these putative enhancers and promoters (Fig. 3a and Supplementary Fig. 3a), suggesting that Prdm1a directly controls these genes. To test these interactions, we cloned putative regulatory elements of the *prdm1a* target *s100a1* into a construct driving *EGFP* (Fig. 3a). In wildtype embryos, the reporter construct for regulatory elements of *s100a1* drives *EGFP* strongly in ear hair cells, but not lateral line hair cells (Figs. 3b-3d), recapitulating endogenous ear expression and demonstrating that these regulatory elements are ear hair cell-specific. We also injected the *s100a1* reporter construct into *prdm1a* mutant embryos and observed ectopic expression in mutant lateral line hair cells (Fig. 3e). These results show that Prdm1a interacts with the regulatory elements of at least some of its putative target genes, and that loss of Prdm1a binding to these elements in reporter constructs leads to ectopic expression in lateral line cells, recapitulating the de-repression of the genes in the mutants.

**Fig. 3.**
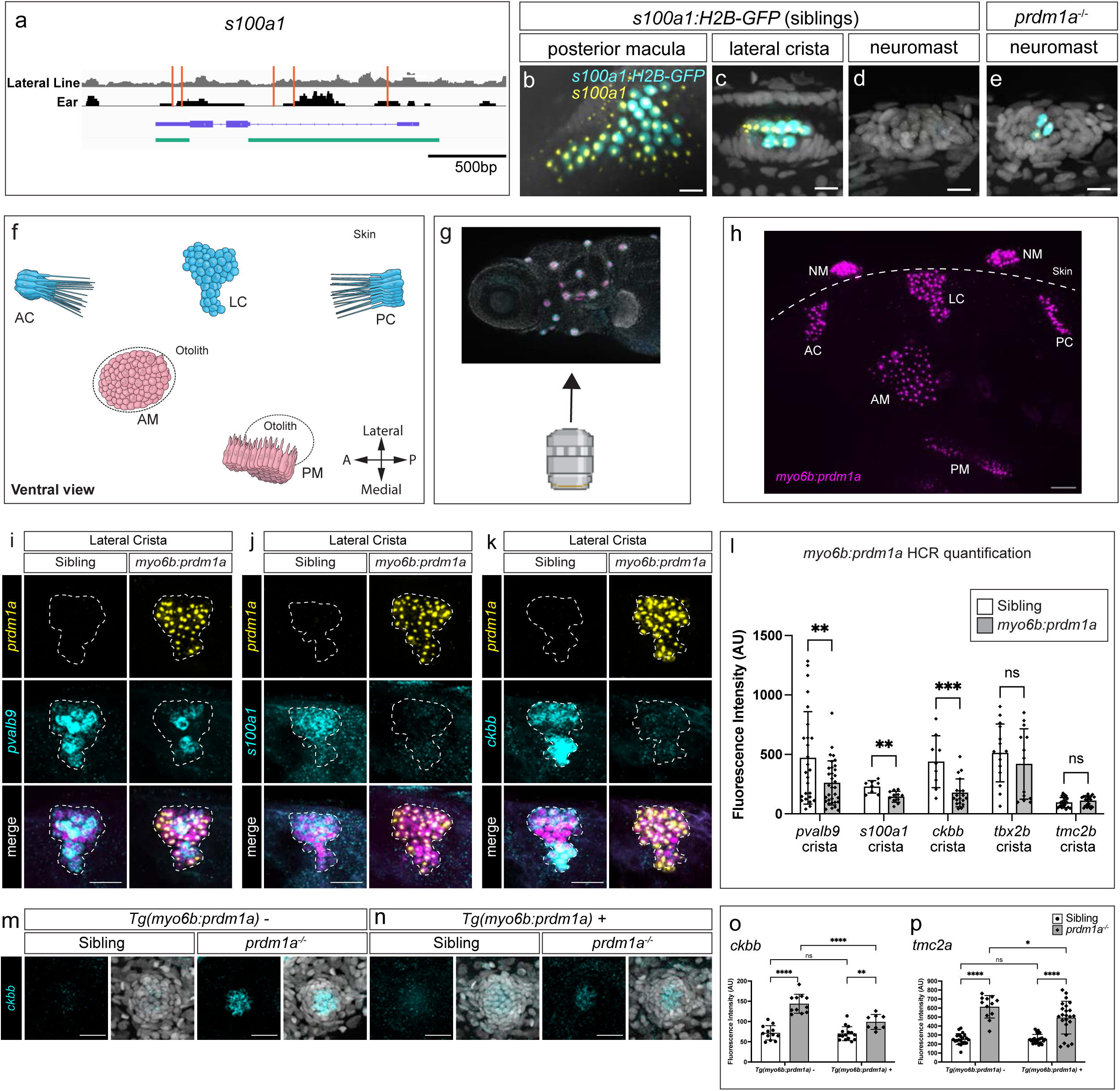
*prdm1a* regulates the expression of target genes in the ear. (a) Schematic of the *s100a1* locus showing lateral line (gray) or ear (black) ATAC-seq reads with location of Prdm1 binding motifs in red. Exons and introns in purple. Cloned enhancer-promoter fusion of *Tg(s100a1:H2B-EGFP)* in green. (b-e) Representative images of *s100a1* HCR (yellow) and H2B-GFP expression (cyan) in 5 dpf wild type embryos driven by *Tg(s100a1:H2B-EGFP)* in the posterior macula (b) and lateral crista (c). 5 dpf sibling neuromast (d) and *prdm1a^-/-^* dorsal neuromast (e). Scale bars= 10 µm. (f) Illustration of the zebrafish ear from a ventral view and orientation of the larva during imaging. AC: anterior crista, LC: lateral crista, PC: posterior crista, AM: anterior macula, PM: posterior macula, NM: neuromast. (g, h) Ventral view of the zebrafish ear, HCR with a *prdm1a* probe (magenta) in a *Tg(myo6b*:*prdm1a)* larva. (i-k) HCR signals for *prdm1a* (yellow), *pvalb9*, *s100a1*, and *ckbb* (cyan) in the lateral crista of sibling and the *Tg(myo6:prdm1a; cryaa:mTurqoise2)* embryos, respectively. (l) Quantification of fluorescent signal from HCRs of *pvalb9, s100a1, ckbb, tbx2* and *tmc2b*. n=8 embryos (two ears each) imaged per probe and genotype. Multiple unpaired t-test, ** p<0.01, *** p<0.001. (m) HCRs of *ckbb* in sibling and *prdm1a* mutant neuromasts. (n) Expression of *ckbb* in *prdm1a^-/-^*hair cells using the *Tg(myo6:prdm1a; cryaa:mTurqoise2)* line downregulates the expression of *ckbb*. (o) Quantification of *ckbb* expression in the various conditions. (p) Quantification of *tmc2a* HCR fluorescence intensity in the various conditions. Two-way Anova with Tukey’s multiple comparisons test, * p<0.05 ** p<0.01, *** p<0.001, **** p<0.0001.

We also investigated the ability of Prdm1a to regulate its putative target genes by ectopically expressing it in all ear hair cells using a *myo6b:prdm1a* driver (Figs. 3i-3k). We performed HCRs for three genes upregulated in mutant lateral line hair cells and quantified expression in the lateral cristae of the ear in ectopic *prdm1a-*expressing larvae and siblings. The cristae were imaged from a ventral view (Figs. 3f-3h). The putative *prdm1a* target genes *pvalb9, s100a1* and *ckbb* are repressed in the ear in the presence of ectopic *prdm1a* as indicated by reduced HCR signal intensity (Figs. 3i-3l). *tbx2b*, the ortholog of mammalian Tbx2, is an essential transcription factor for hair cell fate and development in ear hair cells,^14, 77^ and is expressed in ear hair cells and support cells (Supplementary Fig. 2n). After ectopic expression of *prdm1a* in ear hair cells, *tbx2b* is not significantly repressed in hair cells. As *tmc2b* is expressed in both sibling and *prdm1a* mutant lateral line hair cells (Supplementary Fig. 2l) it served as a control and was indeed not affected by the ectopic expression of *prdm1a* in the ear (Fig. 3l). We also tested if overexpression of *prdm1a* represses expression of selected ectopically expressed ear genes in *prdm1a* mutant hair cells and indeed observed significant downregulation of *ckbb* and *tmc2a* (Figs. 3m-3p).

Taken together, the de-repression of *prdm1a* targets in the lateral line, the repression of these targets upon ectopic *prdm1a* expression in the ear, and the activity of *prdm1a* target gene enhancers all demonstrate a direct genetic interaction between *prdm1a* and at least some of its targets and highlight the central role that *prdm1a* plays in hair cell fate.

### *prdm1a* mutant hair cells have structural changes concordant with expression changes

Different zebrafish hair cell populations exhibit different physical characteristics that are molecularly determined, including length of kinocilia.^16, 78, 79, 80^ Microtubule-based kinocilia are components of the mechanosensory apparatus at the apex of the hair cell. To test if the *prdm1a* mutation impacts these features, we used a *pou4f3:gap43-EGFP* transgenic line and measured kinocilia length in sibling and mutant lateral line hair cells. Sibling kinocilia are 23 µm in length and mutant kinocilia are on average 12 µm long. Thus, *prdm1a* mutant hair cells show a significant reduction in kinocilia length (Figs. 4a-4c). Macula and cristae in the ear have kinocilia lengths of 8 µm and 30 µm, respectively.^16, 78, 79^ Therefore, a shortening of lateral line kinocilia resembles a macular hair cell morphology, as macula hair cells possess the shortest kinocilia of zebrafish hair cell populations. We also measured kinocilia lengths of hair cells of the ear lateral crista in the *myo6b:prdm1a* larvae, and observed a reduction in length from 30 µm to 23 µm in the presence of ectopic *prdm1a* (Figs. 4d and 4e). This shortening of kinocilia is concordant with the repression of *prdm1a* target genes observed in the same transgenic embryos (Figs. 3i-3l), and a partial fate shift toward lateral line characteristics of shorter kinocilia. Mutant lateral line hair cells also possess a higher cell volume (Figs. 4f and 4g) and an enlarged cuticular plate area (Figs. 4h-4iG). However, hair cell polarity is not affected (Figs. 4h and 4j). Mutant lateral line hair cells are likely functional as they take up Daspei and they can be killed by neomycin, which requires functional mechanosensory channels (Figs. 4k-4l).

**Fig. 4.**
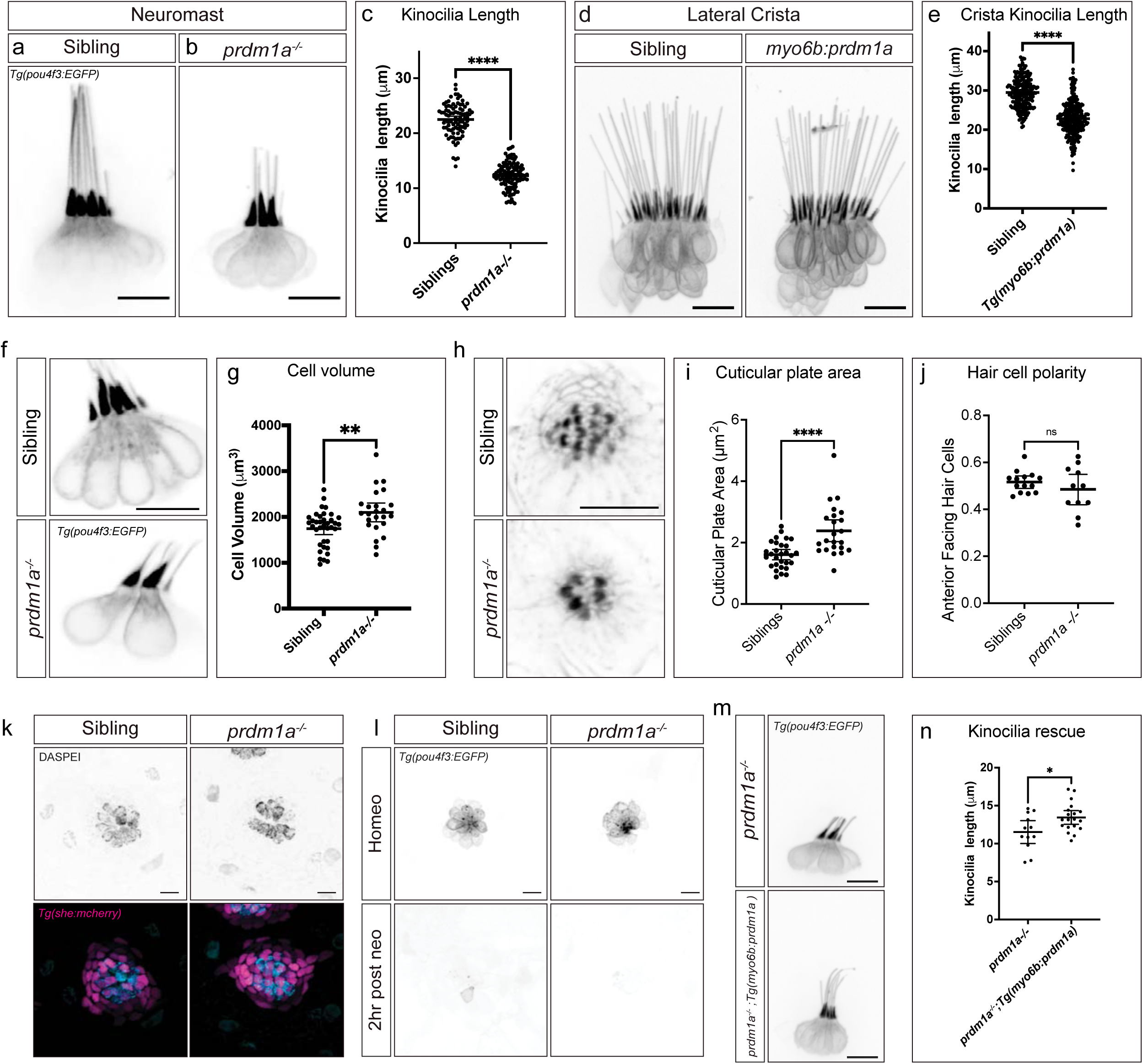
*prdm1a* mutant hair cell morphology changes concordant with expression changes. Fig. 4. (a,b) *Tg(pou4f3:EGFP)* in sibling and *prdm1a^-/-^* neuromast hair cells. (c) Quantification of neuromast kinocilia length in sibling and *prdm1a^-/-^* hair cells. n=11 sibling neuromasts, 89 kinocilia; n=21 *prdm1a^-/-^* neuromasts, 117 kinocilia. Student’s t-test, p<0.0001. (d) *Tg(pou4f3:EGFP)* expression in sibling and *Tg(myo6b*:*prdm1a)* cristae. Scale bars= 10 µm. (e) Quantification of *Tg(myo6b*:*prdm1a)* cristae kinocilia length. n= 15 sibling cristae, 226 kinocilia; n= 15 *Tg(myo6b*:*prdm1a)* cristae, 258 kinocilia. Student’s t-test, p< 0.0001. (f) *Tg(pou4f3:EGFP)* labeling cell bodies in sibling and *prdm1a^-/-^* lateral line hair cells. (g) Quantification of cell body volume in sibling and *prdm1a^-/-^* lateral line hair cells. n=12 sibling neuromasts, 37 cell bodies; n=11 *prdm1a^-/-^* neuromasts, 24 cell bodies. Student’s t-test, p=0.0026. (h) Phalloidin staining of sibling and *prdm1a^-/-^* neuromasts. (i) Quantification of cuticular plate area in sibling and *prdm1a^-/-^* neuromasts. n=14 sibling neuromasts, 29 cuticular plates; n=10 *prdm1a^-/-^* neuromasts, 23 cuticular plates. Student’s t-test p<0.0001. (j) Quantification of the percentage of anterior-polarized hair cells per neuromast. n=14 sibling neuromasts and n=10 *prdm1a^-/-^* neuromasts. Student’s t-test p=0.2916. (k) DASPEI uptake in sibling and *prdm1a^-/-^* lateral line hair cells. (l) *Tg(pou4f3:EGFP)* in sibling and *prdm1a^-/-^* lateral line hair cells during homeostasis and 2 hours after neomycin treatment. (m) *Tg(myo6b:prdm1a)* overexpression in *prdm1a^-/-^* hair cells. (n) Quantification of kinocilia length in *Tg(myo6b:prdm1a)* positive and negative *prdm1a^-/-^* hair cells. n=12 *Tg(myo6b:prdm1a)* negative neuromasts, 12 kinocilia; n=19 *Tg(myo6b:prdm1a)* positive neuromasts. Student’s t-test p=0.0174. All Scale bars= 10 µm.

As *prdm1a* is also expressed in support cells during early stages of regeneration, we wondered in the hair cell defect is possibly non-cell autonomous. However, overexpression of *prdm1a* with the hair cell-specific *myo6b*-promoter increases the kinocilia length, demonstrating that the kinocilium defect in *prdm1a* mutant hair cells is cell-autonomous (Figs. 4m and 4n).

Taken together, the changes in kinocilia length when manipulating *prdm1a* expression in different hair cell types provide evidence that the transcriptional changes we observe correlate with physical differences in hair cells, reinforcing the idea of *prdm1a* controlling hair cell fate in the zebrafish.

### Integrated scRNA-seq analyses suggest *prdm1a* mutant hair cell fate switch to striolar hair cells

While our data provide convincing evidence that *prdm1a* is central to zebrafish lateral line hair cell fate, we asked how complete the conversion to an ear hair cell fate is in the absence of *prdm1a*. We therefore integrated our lateral line scRNA-seq data with two published scRNA-seq studies of zebrafish ear hair cells. Specifically, we integrated an otic subset from a whole-embryo developmental atlas and an ear specific dataset,^5, 75^ which we term “embryo otic subset” and “ear scRNA-seq” respectively (Figs. 5a-5e, Supplementary Data 3). We observed clustering of the *prdm1a* mutant lateral line hair cells with wildtype ear hair cells from both ear datasets, indicating that *prdm1a* mutant lateral line hair cells share transcriptional similarity with ear hair cells (Figs. 5e, Supplemental Data 4). When we label the ear hair cell clusters by annotated cell type,^5^ the *prdm1a* mutant hair cells overlap most with striolar hair cells of the macula (Fig. 5f, Supplementary Data 4), consistent with the shortening of kinocilia. In addition, a differential gene expression analysis comparing *prdm1a* mutant lateral line, sibling lateral line, and ear hair cells shows that a large group of genes is shared between the mutant lateral line and ear hair cells, but not sibling lateral line (Fig. 5g, Supplementary Data 5). While the fate switch is not complete based on transcriptional changes, as some lateral line specific hair cell genes in the *prdm1a* mutant hair cells are not completely downregulated (Supplementary Data 4 and 5), the fate change is nevertheless dramatic.

**Fig. 5.**
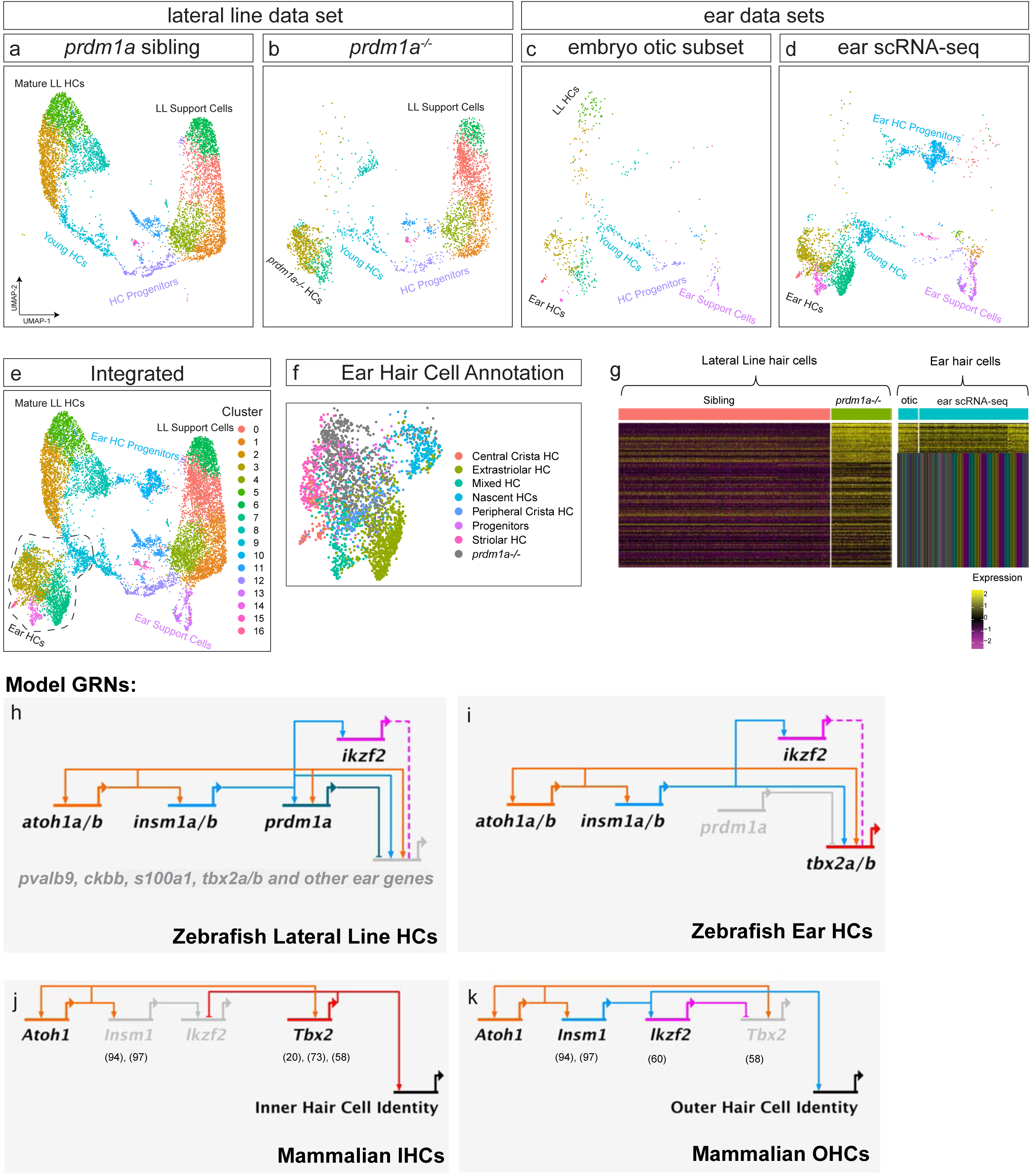
Integrating ear scRNA-seq to build a hair cell differentiation GRN. (a-e) UMAPs of integrated scRNA-seq, showing sibling lateral line, *prdm1a^-/-^*lateral line, embryo otic subset, ear scRNA-seq, and complete integration respectively. (f) Integrated ear *prdm1a^-/-^* hair cell cluster with cells color coded by cell type from the ear scRNA-seq annotation.^5^ *prdm1a^-/-^* hair cells are in grey. (g) Heatmap of mature hair cell genes of the triple integrated scRNA-seq data set divided into *prdm1a^-/-^* lateral line cells, ear hair cells of the embryo otic subset and ear scRNA-seq data sets. (h) GRN of zebrafish lateral line hair cell specification and development. (i) GRN of zebrafish ear hair cell specification and development. (j) GRN of mouse inner hair cell specification and development. (k) GRN of mouse outer hair cell specification and development. Numbers in brackets indicate citations for the individual interactions.

Here we show that within the lateral line, *prdm1a* promotes proliferation of support cells and the development and regeneration of lateral line hair cells. *prdm1a* also suppresses an ear hair cell fate through its role as a transcriptional repressor. Loss of *prdm1a* in the lateral line leads to de-repression of a large group of ear-specific hair cell genes, resulting in physical changes in lateral line hair cells consistent with an ear macula hair cell identity. We provide multiple lines of evidence – through scRNA-seq, ATAC-seq, reporters of regulatory element activities, and ectopic *prdm1a* expression experiments – that *prdm1a* represses ear fate in lateral line hair cells. Importantly, our integrated scRNA-seq analysis of *prdm1a* mutant and wild type ear hair cells shows that the fate conversion is almost complete. Therefore, the expression of *prdm1a* during hair cell maturation reflects an inflection point in the determination of hair cell fate between lateral line and ear. The differential regulation of *prdm1a* in the lateral line and the ear likely played an important role in the evolution of these two hair cell types and also demonstrates that individual transcription factors play key roles in determining hair cell subtypes also during regeneration.

## Discussion

Questions of how individual genes and gene regulatory networks determine cell fate are central to developmental biology, and foundational works in the field have broad influence over research to this day.^81, 82, 83, 84, 85^ We investigated the role of *prdm1a*, central to cell fate decisions, with wide implications for linking evolution and specification of cell fate with regenerative genetics to restore hearing. We uncovered two roles for *prdm1a* in the lateral line: determining hair cell fate and promoting proliferation during development and regeneration.

We focused on the significant role of *prdm1a* in determining hair cell fate. Our integration of *prdm1a* mutant scRNA-seq data with two zebrafish ear hair cell scRNA-seq datasets shows how a single transcription factor can shift cell fate with consequences for transcriptional identity and physical characteristics that affect cell function. When comparing all three zebrafish hair cell types (sibling, *prdm1a* mutant lateral line, and the wildtype ear), it is important to note that although there are transcriptional and morphological differences between the three populations, they are all hair cells. All three hair cell types share genes along the differentiation trajectories, with more genes in common than different (Supplementary Data 3 and 4). Therefore, the three datasets present an ideal foundation from which to build GRNs describing hair cell development and fate.

Just as there are transcriptionally and morphologically distinct hair cells in the zebrafish ear and lateral line,^5, 79^ there are also unique hair cell types in the mammalian cochlea and vestibular organs.^86^ In both the ear and lateral line, the genes that initiate specification and differentiation of hair cell progenitors have been well characterized.^87, 88, 89, 90, 91, 92, 93^ Specifically, *Atoh1* and *Insm1* are key transcription factors for hair cell specification and differentiation across species, and their epistatic interactions have been investigated in mice.^94, 95^ By using our ATAC-seq data to identify conserved transcription factor binding sites and incorporating the developmental timing of expression, we infer the existence of a coherent feed forward subcircuit^96^ in the lateral line involving *atoh1a/b* and *insm1a/b* activating downstream hair cell genes, including *prdm1a* (Figs. 5h and 5i, Supplementary Fig. 3b). The presence or absence of *prdm1a* in this subcircuit determines lateral line or ear hair cell fate.

As *prdm1a* is the key gene in determining ear versus lateral line hair cell fate in the zebrafish, we compared its place in the GRN to another hair cell fate specification process; the inner-versus outer hair cell fate decision in the mammalian cochlea. The network of *Insm1, Ikzf2,* and *Tbx2* in cochlear hair cells determines if they will develop into inner or outer hair cells, each with different morphologies and functions in hearing.^14, 58, 60, 77, 94, 97^ While these studies each uncovered key aspects of this fate decision, a unified network has not yet been constructed. We built a GRN underlying the inner-, versus outer hair cell fate decision in the mouse cochlea (Figs. 5j and 5k) based on published data and compared it to the model zebrafish GRNs for the lateral line and ear (Figs. 5h and 5i).

The zebrafish GRNs possess striking similarities to the mouse ear hair cell fate GRN. The same coherent feedforward loop with *Atoh1* and *Insm1* appears to be present in both models, and the expression of either *Tbx2* or *Ikzf2* determines the fate of developing hair cells.^60, 94^ This similarity between hair cell GRNs in the zebrafish ear, and mouse ear, and lateral line reveals a shared architecture of the hair cell determination and differentiation network implying deeply conserved systems at the heart of hair cell development and fate.

The expression of *prdm1a* is a key difference between lateral line-, and ear hair cells. However, *prdm1a* and *Prdm1* are both expressed in embryonic developing zebrafish ears (Supplementary Figs. 1b and 1c) and mammalian ears, respectively, ^98^ but repressed in sensory epithelia and completely eliminated from ears by the time hair cells mature and function. We therefore suggest that ancestral hair cells diverged into ear and lateral line fates based on the differential regulation of *prdm1a*.

It is well understood that transcriptional repression of a GRN branch or node is a powerful tool to direct cell type specification in developing tissues.^99, 100, 101^ The repeated deployment of *Prdm1*/*prdm1a* in cell fate decisions in hematopoiesis and photoreceptors in mice,^45, 46, 47^ neural crest-, muscle-, and hair cells in zebrafish^48, 49, 50^ highlights the potency of its activity as a transcriptional regulator across tissues and species. In all these cases, the mutation of *Prdm1*/*prmd1a* leads to a de-repression of a large number of genes and shift in cell fate. Our study cements *prdm1a* in this role as a master regulator of cell fate and the possible conservation of its function across evolution. The results not only inform our understanding of how hair cell types might have evolved but also serves as a template to induce correct hair cell differentiation via the manipulation of key transcription factors during regeneration in the cochlea.

## Methods

### Zebrafish

Work with zebrafish followed standard procedures and was overseen by the IACUC at the Stowers Institute for Medical Research. The following transgenic lines were used: *Tg(she:H2A-mCherry)^psi56Tg^, Tg(she*:*H2B-EGFP)^psi59Tg^, Tg(myo6:H2B-mScarlet-I)^psi66Tg^*^102^, *Tg(pou4f3:Gap43-EGFP)*^103^ and *Tg(sqet20Et:EGFP)*^104^. The *prdm1a^psi91^* mutant was identified in an ENU-induced mutagenesis screen.

### Confocal Microscopy

Embryos were embedded in 0.8% low melting point agarose (Promega) and mounted on 35mm glass bottom dishes (Mat Tek). Imaging was performed on a Nikon Ti2 Yokogawa CSU-W1 spinning disk microscope with a Hamamatsu Orca Fusion sCMOS camera. Images were acquired with either 10x CFI Apo LWD or 40x WI 1.15 NA Lambda Objectives. Images were processed using Imaris versions 8.3.1 and 9.6.1 (Bitplane) and Fiji ^105^.

### HCR In Situ Hybridization

All HCRs were performed using probes and reagents from Molecular Instruments according to standard zebrafish embryo protocols,^106^ with the following changes: Embryos were permeabilized in 80% acetone at -20°C for 30 minutes, followed by washing in PBST 3x10 minutes prior to probe pre-hybridization. All reaction volumes were reduced by half, but probe and amplifier concentrations maintained with protocol recommendations; amplification step was extended to 48 hours. All probes were custom synthesized by Molecular Instruments.

### HCR Signal Quantification

The HCR signal was normalized with a 50μm rolling ball background subtraction in Fiji. The signal was then quantified by drawing ROIs around cells of interest, and using the Measure function to measure mean signal intensity.

### Kinocilia length quantification

Using images of *Tg(pou4f3:EGFP)*, maximum intensity projections of z-stacks were created in Fiji and lengths of individual kinocilia measured using the Measure function.

### Cuticular plate measurements

Cuticular plate measurements were obtained by manually drawing ROIs around the phalloidin signal of stained cuticular plates and using the measure function in Fiji.

### Daspei staining

To visualize hair cells embryos were placed in a 0.006% solution of DASPEI (2-(4- (dimethylamino)steryl)-N-ethylpyridinium iodide, [Invitrogen, USA]) diluted in embryo media for 10 minutes. Embryos were then washed and anesthetized with MS-222 for imaging on a spinning disk microscope.

### Phalloidin staining

Phalloidin staining was performed using Alexa Fluor 488-conjugated phalloidin (Thermo Fisher Scientific). Zebrafish were fixed in 4% paraformaldehyde with 1% PBS for 4 hours at 4□°C, then washed in PBS containing 1% DMSO, 0.5% Triton X-100, and 0.1% Tween-20 (PBDTT buffer). Phalloidin (1:40, #A12379, Thermo Fisher Scientific) and DAPI (1:1000) were then added in PBDTT buffer supplemented with 1% BSA and incubated at room temperature for 2 hours.

### Cell Body Measurements

To measure hair cell body volumes, masks were drawn manually around the body of hair cells, excluding the kinocilia, in Fiji. Using the measure function in Fiji, the hair cell body volume was estimated as an ellipsoid from the major and minor axes of the hand drawn ROI output by Fiji.

### Hair Cell Ablation

Lateral line sensory hair cells were ablated in 5 dpf larvae by incubation in 300μM neomycin (Sigma Aldrich) in standard embryo media for 30 minutes at 28°C.^52^ Neomycin was washed out thoroughly and embryos returned to 0.5x E2 embryo media until analysis timepoint.

### Cell Proliferation Assay

Proliferating cells were assayed as previously described.^107^ 5 dpf larvae were incubated in 3.3mM EdU (Carbosynth) in 1% DMSO in standard embryo medium for 48 hours following hair cell ablation. Larvae were then fixed in 4% paraformaldehyde overnight at 4°C. All subsequent steps were performed at room temperature. Larvae were washed 3x10 minutes in PBS/0.8% Trition-X (PBSTX), blocked in 3% BSA in PBSTX for one hour, and washed 3x5 minutes in PBS. To stain, embryos were incubated in staining cocktail consisting of 1x TBS, 2mM CuSO_4_, 2.5uM Alexa-647-Azide (Invitrogen), 50mM ascorbic acid, and 0.4% Triton-x for 30 minutes. After staining, larvae were washed 3x10 minutes in PBSTX, stained with 5ug/ml DAPI at RT for five minutes, and imaged as described above.

### scRNA-seq

#### Larva dissociation and FACS

600 each *prdm1a* mutant and sibling embryos with the *Tg(she:H2A-mCherry)^psi56Tg^* lateral line marker were separately dissociated in 4.5ml 0.25% Trypsin-EDTA (Gibco) for 3.5 minutes on ice by triturating with a 1ml pipette repeatedly. Supernatant containing cells was filtered through a 70μm Cup Filcon (BD Biosciences) and cells were pelleted by centrifugation at 720g for five minutes at 4°C. Cells were washed in cold PBST by resuspending the pellet and pelleted again at 720g for five minutes at 4°C. Cells were filtered through a 35μm strainer (Falcon-Corning), and stained with Draq5 (BioStatus) at a dilution of 1:500. Cells were then sorted on a BD Influx Cell Sorter (BD Biosciences). mCherry and Draq5 double positive cells were sorted into 90% MeOH for fixation and stored at 4°C until rehydration.

#### 10x Chromium scRNA-seq library prep

FAC-sorted MeOH fixed cells prepared as above were processed according to 10X Genomics standard protocols. Cells were rehydrated in 1% BSA and 0.5 U/μl RNase-inhibitor in ice-cold DPBS. A volume of 34ul was loaded onto the Chromium Controller, targeting ∼12,000 cells per sample. Library prep was performed using the Chromium Next GEM Single Cell 3’ GEM, Library & Gel Bead Kit v3.1. Two independent replicates were sequenced for both *prdm1a* mutant and sibling conditions. Libraries were pooled and sequenced on an Illumina NextSeq 500 in High Output mode, with 28 bp, 8bp, and 91 bp for read 1, i7 index, and read 2 respectively.

### Single cell RNA-seq Data Analysis

Data were processed using 10X Genomics Cellranger v4.0, first by demultiplexing using cellranger mkfastq with default parameters, then cellranger count was used to align reads to the GRCz11 genome (Ensembl release 98) and generate a feature count matrix. The matrix was then loaded into R package Seurat (v4.3)^108^ for further bioinformatics analysis. First, the top 2000 variable genes were selected using the loess span parameter for principal component analysis. Based on elbow plot, 30 PCs were selected to generate the nearest neighbor graph, clustering as well as the UMAP visualization were done using Seurat default parameters. External scRNA-seq data used for integration were downloaded from GEO. All replicates and samples were then merged and integrated using Seurat’s default CCA method for downstream analysis. For marker gene detection, we used the Seurat FindAllMarkers function (logfc.threshold=0.25) to generate differential expressed genes for every cluster and annotated based on ZFIN database knowledge. Heatmaps were then generated using the DoHeatmap function. All raw and processed sequencing data have been deposited into NCBI Gene Expression Omnibus (GEO) database with accession number GSE268538.

### ATAC-seq

#### Larva dissociation and FACS

ATAC-seq was performed generally following a previously established protocol,^109^ with some modifications. 600-1200 *Tg(she:H2A-mCherry)^psi56Tg^* embryos were fixed in 1% PFA in PBS at 4°C for 30 minutes. Glycine was added to a final concentration of 125mM for five minutes to quench formaldehyde. Embryos were moved to 4.5ml 0.25% Trypsin-EDTA (Gibco) and dissociated for ∼8 minutes on ice by triturating with a 1ml pipette repeatedly. Supernatant containing cells was filtered through a 70μm Cup Filcon (BD Biosciences) and cells were pelleted and washed twice by centrifugation at 720g for five minutes at 4°C and resuspending in cold PBST. Cells was filtered through a 70μm Cup Filcon (BD Biosciences) one additional time and pelleted by centrifugation at 720g for five minutes at 4°C. Cells were resuspended in 4ml PBST and stained with Draq5 at a dilution of 1:500. mCherry and Draq5 double positive cells were then sorted into pools of 50,000 on a BD Melody Cell Sorter (BD Biosciences) into chilled 1.5ml centrifuge tubes.

### Tagmentation

Cells were pelleted by spinning in a microcentrifuge for 5 mins at 4°C and 1000g. Supernatant was removed carefully by pipette, and cells were resuspended in tagmentation reaction mix (22.5 μl H_2_O, 25 μl TD Buffer (Illumina), and 2.5 μl TD Enzyme (Illumina)) and incubated in a ThermoMixer (Eppendorf) at 37°C for 30 minutes, 1000 RPM. 50 μl reverse crosslinking buffer (final concentration of 50mM TRIS pH 8, 1mM EDTA, 1% SDS, 200mM NaCl, 5ng/ml Proteinase K (Roche)) was added and mixture was incubated at 65°C overnight in a ThermoMixer. Tagmented DNA was purified with a DNA Clean & Concentrator 5 kit (Zymo Research) and eluted in 20 μl H2O. Library PCR amplification was performed with 20 μl eluted DNA, 25 μl NEBNext Ultra II Q5 Master Mix, 2.5ul forward primer Ad1 25mM stock, 2.5ul reverse barcode primer^110^ Ad 2.x 25mM stock. PCR conditions as follows: 72°C 5 mins, 98°C 30 seconds, 11 cycles of 98°C 10 seconds, 63°C 30 seconds, 72°C 1 minute, 4°C hold. Library was size selected with SPRIselect beads (Beckman Coulter) with the following conditions: 0.8x volume of beads first selection, the supernatant was retained and an additional 1.0x volume of beads was added (total volume 1.8x). The bead pellet was retained and rinsed with 85% EtOH, dried and DNA was eluted in 30ul H_2_O. Libraries were analyzed on a Bioanalyzer 2100 (Agilent) to evaluate tagmentation and quantified on a Qubit 3.0 (Thermo Fisher).

### Sequencing and Read Alignment

ATAC libraries were pooled and sequenced on an Illumina NextSeq 500, 2x75bp paired-end mode. The zebrafish reference genome and transcriptome was obtained from Ensembl (version: GRCz11-102).^111^ It was segmented into contiguous windows at 50bp resolution using makewindows function of the bedtools package.^112^ These windows were used as a basis for this analyses. Sequencing reads for each sample were mapped to the genome using the STAR aligner (v 2.7.9)^113^ (using parameters –alignIntronMax -1 – alignIntronMin -1 –outFilterMultimapNmax 1 –outFilterMultimapScoreRange 1 – outFilterMismatchNmax 3), producing corresponding BAM files. A coverage matrix was tabulated by counting reads in each sample’s BAM file (column) overlapping each 50bp window (row) using featureCounts.^114^ The coverage matrix was then quantile-normalized across all samples using R package preprocessCore and filtered, removing windows (rows) either overlapping a genome-wide blacklist (as determined in forthcoming research or exhibiting insufficient coverage (specifically, having no group with a mean coverage of at least 20).

### Peak Calling and Putative Regulatory Element Determination

A peak set (in narrowPeak format) was determined separately for each experimental group using Genrich^115^ to analyze the group’s replicate BAM files (with parameters ‘-a 0 -q .05 -g 250 -l 100’ and to ignore regions appearing in the genome wide blacklist). A consensus peak set for the mark as a whole was formed (again in narrowPeak format) by merging overlapping or proximal (within 100bp) peaks from these group-specific peak sets using capabilities provided by BioConductor’s GenomicRanges::reduce.^116^ Candidate regulatory elements were defined as the subset of ATAC-Seq consensus peaks being both (a) overlapping or proximal to (within 100bp) any H3K27ac or H3K4me3 consensus peak (as determined in forthcoming research) and (b) exhibiting a minimum degree of accessibility, by overlapping, at least in part, the filtered coverage matrices. A single Ensembl gene was assigned to each putative regulatory element as its putative target based on proximity, as follows. It was classified as a promoter targeting the closest protein coding gene to which it is within 250bp upstream or 100bp downstream, if any; otherwise, it was classified as an enhancer targeting the closest protein coding gene, considering only those previously found to have been expressed in our lateral line scRNA-seq atlas.^19^

### Motif Searches

Using the list of genes upregulated in *prmd1a* mutant hair cells and ATAC-seq peaks described above, we compiled a list of enhancers and promoters associated with those upregulated genes. We used Samtools^117^ to extract the genomic sequence belonging to the peak list and the MEME suite tool AME^118^ to search for enriched TF motifs, with a background sequence set consisting of shuffled sequences preserving the dinucleotide frequency of the input set. We used the JASPAR 2022 vertebrate core TF motif database^119^ as the reference for AME.

### Generation of Transgenic Reporter Lines

We identified ATAC peaks associated with s100a1 and designed primers to clone the gene’s enhancer and promoter. To clone the 3’UTR peak primers s100a1: F-5’ACTTTCATGTGTTACTTCATTTGAAGGAAAATG’3, R-5’TTTAACCATTTTATTTTGCTTCTGTCAAAATGTACATC’3 were used. To clone the upstream promoter primers F-5’CAGAATTCAGCCAAAACGTCTTAAAGCATGG’3, R-5’GTTAGAAAATATTATCTGCAAGATAAATAATTAAATACAAGTC’3 were used. These regions were then inserted together into a miniTol2 120 plasmid containing a beta-actin minimal promoter upstream of H2B-GFP followed by the SV40 poly adenylation signal via Gibson assembly 121. This reporter construct, Tg(s100a1:H2B-EGFP), was then injected at a final concentration of 30ng/μl with tol2 mRNA into one-cell stage zebrafish embryos for Tol2-mediated integration122, 123. Injected embryos were screened for EGFP expression at 5 dpf and raised for outcrossing with wildtype fish to establish stable transgenic lines. The Tg(myo6:prdm1a; cryaa:mTurqoise2) construct was created using gateway assembly and the Tol2Kit integration122, 123. An existing p5E entry vector with the myo6b enhancer was used124.The prdm1a cDNA was cloned into the pME entry vector, with the SV40 poly-A signal. cDNA was synthesized using RNA extracted from 5dpf zebrafish embryos with TRIzol (Invitrogen), following standard protocols. RNA was precipitated with isopropanol, washed with 75% EtOH, dried, and resuspended in RNAse-free dH2O. cDNA was synthesized using ProtoScript II Reverse Transcriptase (NEB) with 1ug total RNA and oligo d(T)23VN as a primer, following standard protocols. Following reverse transcription, the reaction volume of 20ul was diluted up to 100ul with dH2O. prdm1a cDNA was amplified with the following primers: F-5’ ATCTCAGGCACTTGCAGGAGAAGTC 3’, R-5’ CTAGGTATCCATGGCCTCCTCTGTCTC 3’. The cryaa promoter was cloned into the p3E entry vector driving mTurquoise using cDNA prepared above using the follwing primers: F-5’ GTGGAGACCCCTGATTAATAAAGGGACTTA 3’, R-5’ AATGTCAGACCTGGTAACTCCTTACTGTAA 3’. The final plasmid was assembled by the gateway reaction using all three entry vectors described above and the pDestTol2pA2 destination vector.

### Statistical Analysis

Statistical tests were performed using GraphPad Prism 10 (Version 10.2.3). When comparing data from more than two groups, statistical significance was calculated using one-way ANOVA with Tukey’s post hoc test. Data from two groups were compared using two-tailed unpaired t-test. *p-value*s smaller than 0.05 were considered to be statistically significant. Statistical details of experiments, such as *p-value*s and sample size are specified individually for each experiment in the Fig. legends. Plots were made in GraphPad Prism 10.

## Acknowledgements

We thank Jillian Blanck, Kevin Ferro, KeyongMin Bae, Fang Liu, and Jeffery Haug for technical assistance and advice performing FAC-sorting. We thank Michael Peterson, Kate Hall, and Anoja Perera for technical assistance and advice in sequencing libraries on NGS instruments. We thank Mark Miller for graphic design. We thank Tatjana Sauka-Spengler and Robb Krumlauf for thoughtful discussions and Piotrowski Lab members Daniela Münch, Julia Peloggia and Aurelie Hintermann for thorough readings of the manuscript. This research was supported by NIH (NIDCD) award 1R01DC015488-01A1 to TP, NIH (NIDCD) award 1F32DC019023-01A1 to JES, funding from the Hearing Health Foundation to TP, and institutional support from the Stowers Institute for Medical Research to TP.

## Author contributions

JES, Y-YT and TP conceived the project and designed experiments. JES, Y-YT, LS, MEL, EE, ARS, and JSK performed experiments. JES, Y-YT, LS and MEL analyzed experimental data. SC, NTTT, and MC, performed computational and biostatistical analysis. JES, Y-YT and TP wrote the manuscript with input from all authors. AS and JF provided the data for otic subset of zebrafish scRNA-seq analysis.

---

**Supplementary Fig. 1.**
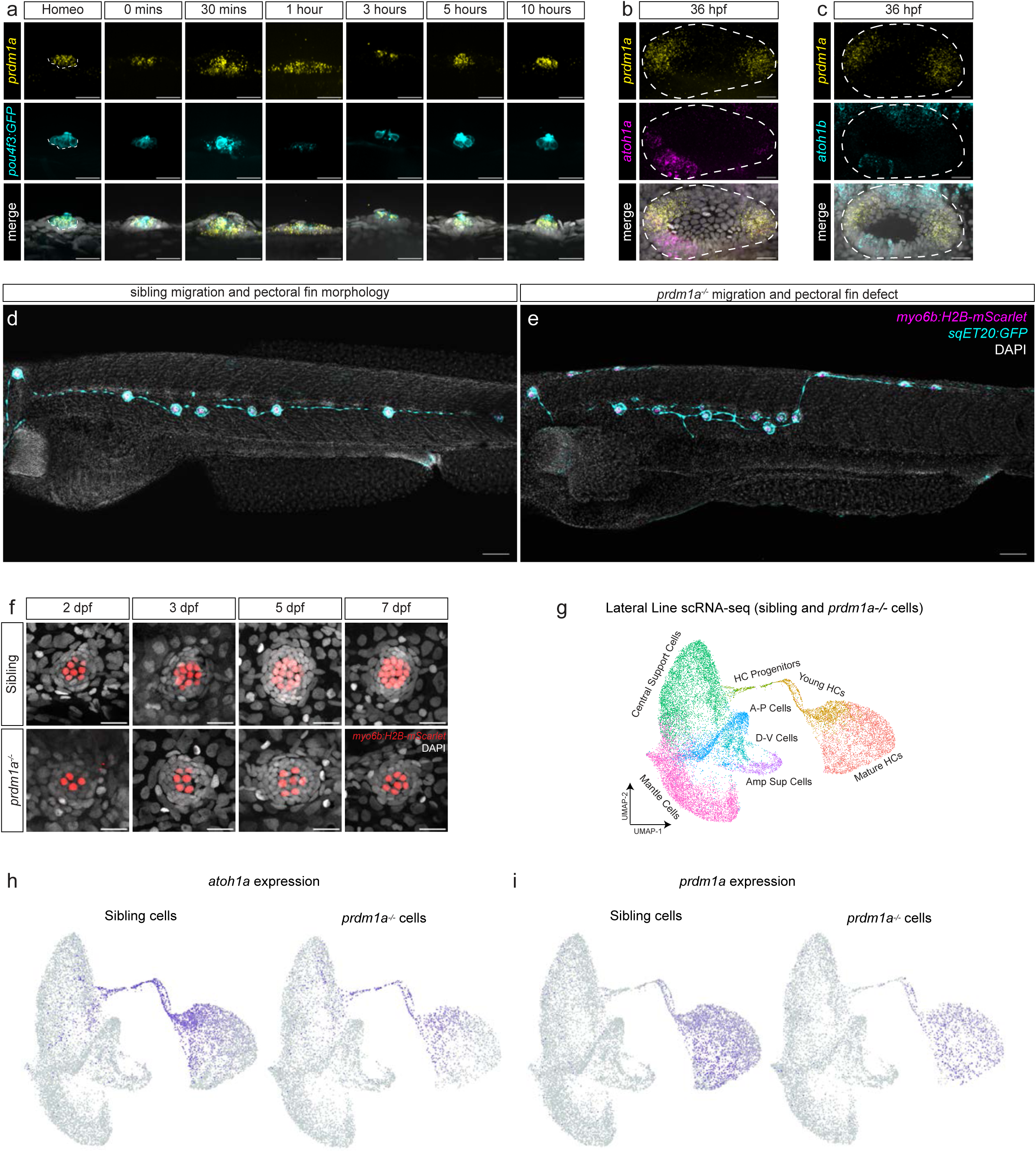
(a) Full regeneration time HCR course of *prdm1a* (yellow) expression in the lateral line with *Tg(pou4f3:GAP-EGFP)* expressed in hair cells. Scale bars= 20 µm. (b) Zebrafish otic vesicle at 36 hours post fertilization with HCR probes for *prdm1a* in yellow and *atoh1a* in magenta. Scale bars= 20 µm. (c) Zebrafish otic vesicle at 36 hours post fertilization with HCR probes for *prdm1a* in yellow and *atoh1b* in cyan. Scale bars= 20 µm. (d, e) Distribution of neuromasts in sibling and *prdm1a^-/-^* embryos respectively, with *Tg(sqET20-EGFP)* marking mantle cells and interneuromast cells and *Tg(myo6b:H2B-mScarlet)* in hair cells. Scale bars= 50 µm. (f) Hair cell developmental time course in *prdm1a^-/-^* and siblings with *Tg(myo6b:H2B-mScarlet)* in hair cells. Scale bars= 20 µm. (g) UMAP of scRNA-seq from sorted lateral line sibling and mutant cells, colored by cell type^19^. When plotted with all cell types present Mantle, A-P, D/V and Amplifying support cells do not express *prdm1a* (H, I). Feature plots of the expression of *atoh1a* and *prdm1a* in the sibling and *prdm1a^-/-^* lateral line.

**Supplementary Fig. 2.**
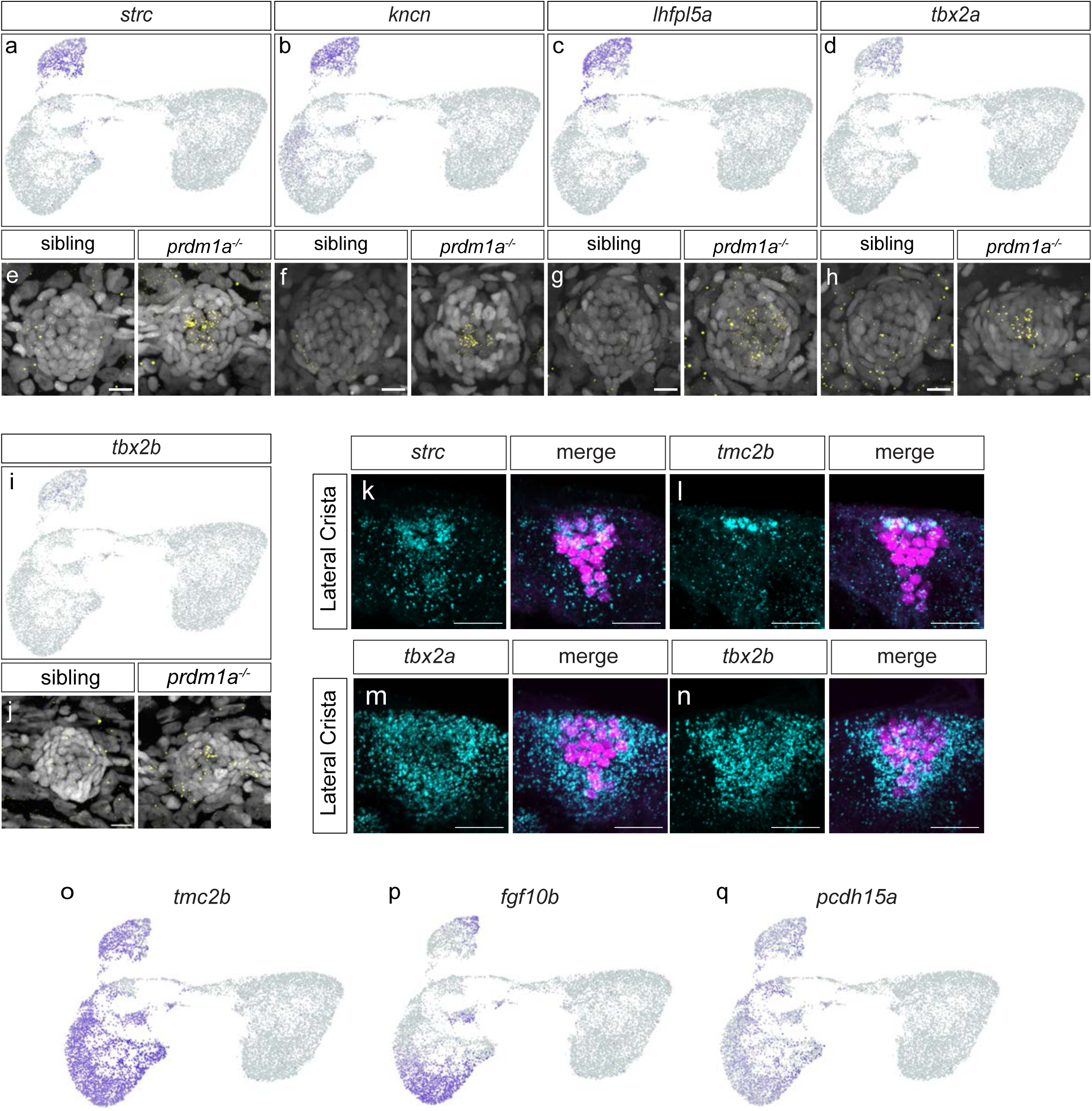
(a-d, and i) Feature plots for *strc, kncn, lhfpl5a, tbx2a* and *tbx2b* respectively in the hair cell developmental trajectory scRNA-seq subset. (e-h, and j) HCRs (yellow) for *strc, kncn, lhfpl5a, tbx2a* and *tbx2b* respectively in siblings and *prdm1a^-/-^ embryos*. Scale bars= 20 μm. (k-n) HCRs (cyan) of *strc, tmc2b, tbx2a and tbx2b* in the lateral crista of the ear in *Tg(myo6b:H2B-mScarlet)* transgenic lavae in which hair cells are labeled in magenta. Scale bars= 10 µm. (o) Feature plot of *tmc2b* that is expressed in sibling and mutant hair cells and is not a *prdm1a* target (p,q) Feature plots of *fgf10b* and *pcdh15a* that are expressed in mature hair cells.

**Supplementary Fig. 3.**
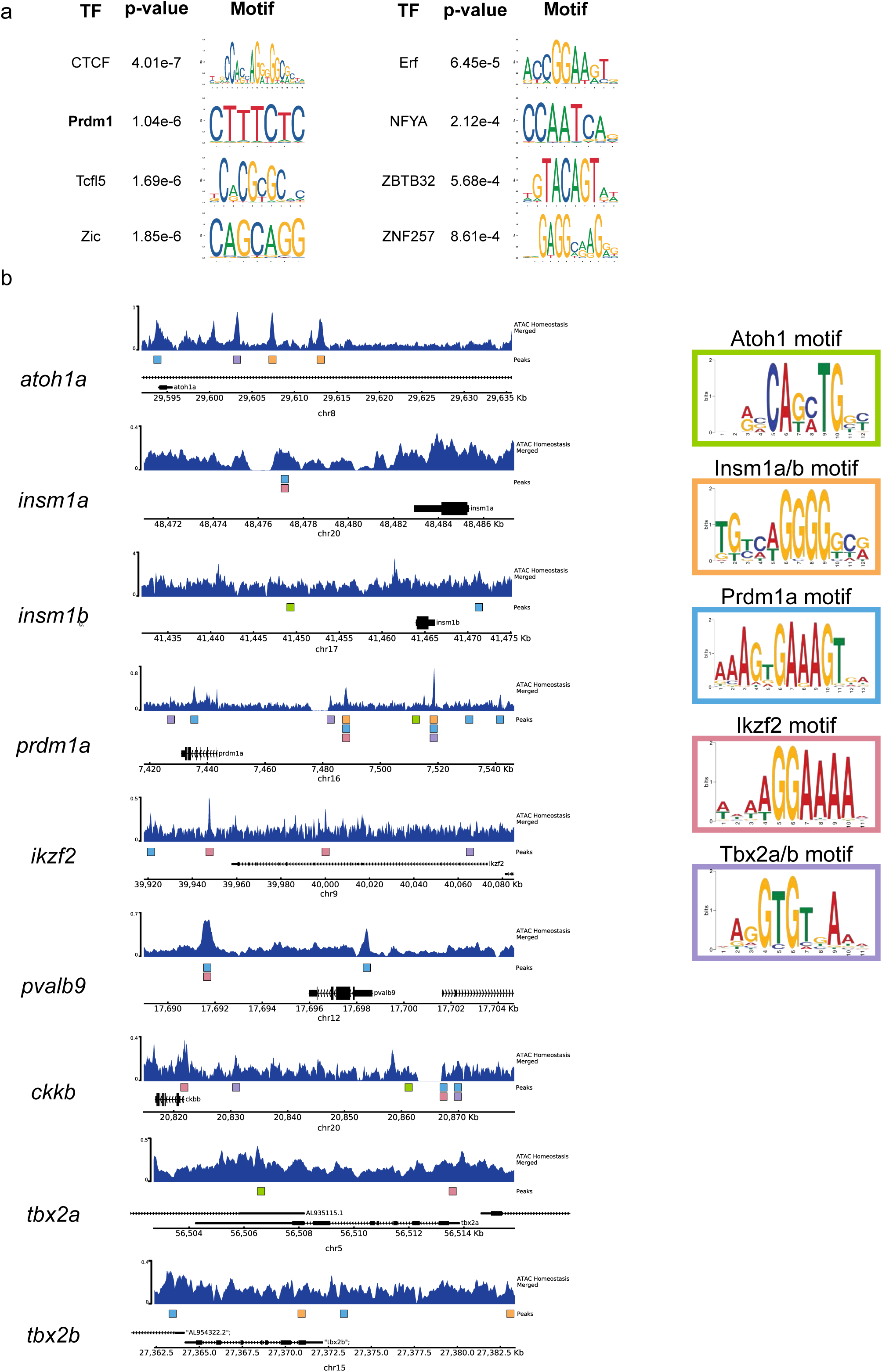
(a) Transcription factor motif enrichment analysis from promoters and enhancers associated with *prdm1a* target genes. (b) ATAC-Seq tracks of FAC-sorted lateral line cells with transcription factor motifs indicated with colored bars. The presence of motifs for putative upstream regulators supports the GRNs proposed in Fig. 5H and 5I.

**Supplementary Table 1:**
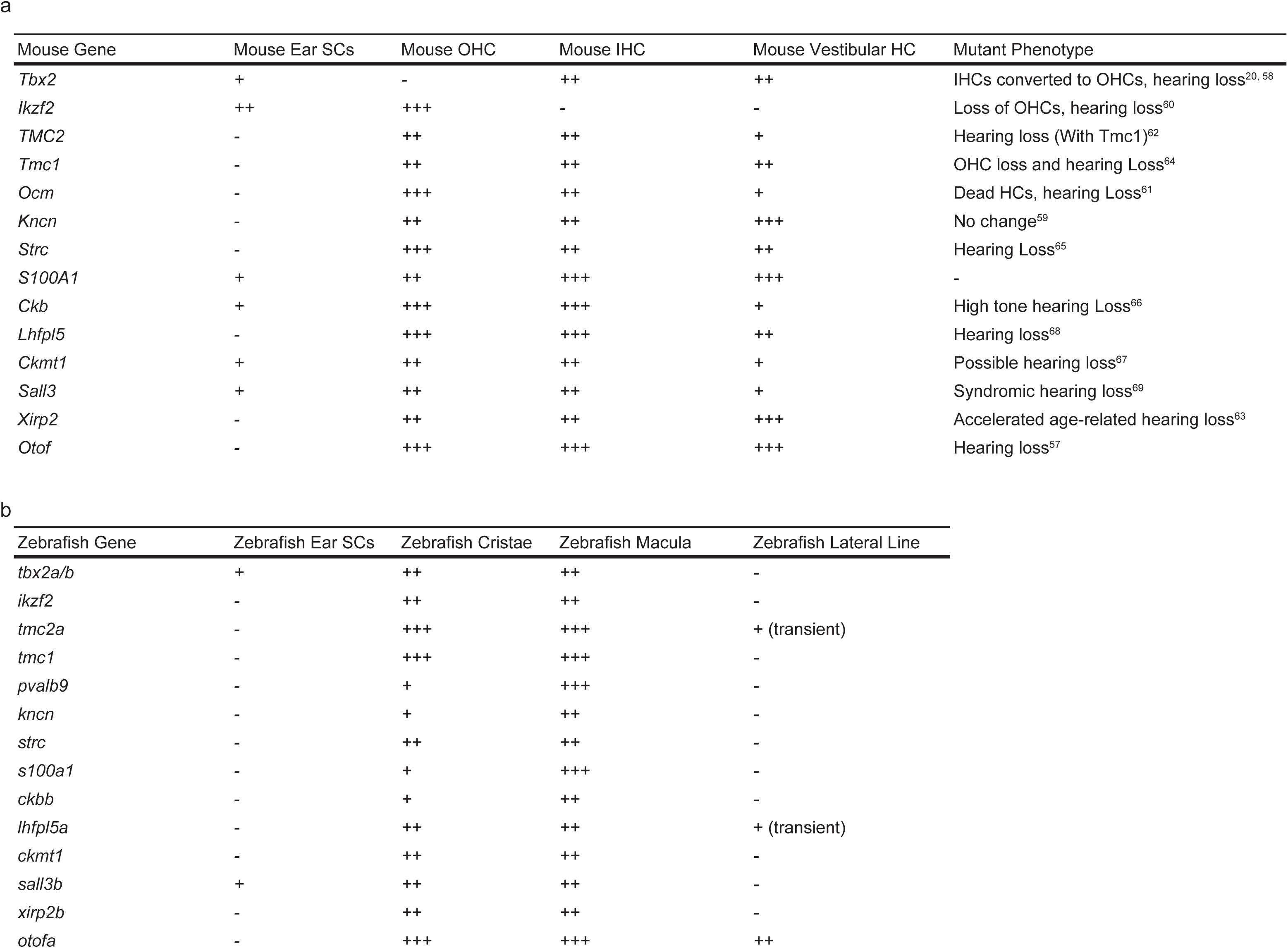
Table of zebrafish and mouse expression of genes upregulated in the *prdm1a^-/-^* lateral line and mutant phenotypes from external data sources. Relates to Fig. 2.

